# CanASM: A comprehensive database for genome-wide allele-specific DNA methylation identification and annotation in cancer

**DOI:** 10.1101/2025.04.03.646980

**Authors:** Jianmei zhao, Zeyu zhao, Hongfei Li, Haojie Yu, Hao Lin, Hanqi Chen, Xuecang Li, Di Liu, Yiming Wang, Guohua Wang

## Abstract

Allele-specific DNA methylation (ASM) provides critical insights into the complex genetic and epigenetic mechanisms regulating gene transcription. Emerging evidence suggests that ASM is particularly enriched in gene enhancer regions, and recent studies have demonstrated that ASM is increased in cancer tissues compared with normal tissues. Despite the increasing recognition of ASM as a potential biomarker in tumorigenesis, systematic resources dedicated to identifying and annotating ASMs in cancer contexts remain limited. In this study, we developed CanASM (https://bioinfor.nefu.edu.cn/CanASM/), the first comprehensive database specifically designed to identify and annotate ASM in cancer. In CanASM, ASM sites identified from bisulfite sequencing data across 31 cancer types and their matched normal tissue samples are cataloged. Importantly, CanASM includes extensive regulatory annotations for ASMs, including associated genes, cis-regulatory elements and transcription factor binding colocalizations, transcription factor affinity changes, etc. Users can query and explore ASMs using various parameters, such as single-nucleotide variations (SNVs), chromosomal coordinates, and gene names. The current version of CanASM includes 5,003,877 unique SNV–CpG pairs, including 3,056,776 index SNVs, of which 2,634,406 are single-nucleotide polymorphisms (SNPs), and 4,157,508 CpGs. With an intuitive interface for browsing, querying, analyzing, and downloading, CanASM serves as a valuable resource for researchers investigating cancer-associated genetic variations and epigenetic regulation in cancer.

**Author summary:** Identifying target genes and regulatory mechanisms of cancer-associated genetic variants remains a major challenge in the post-GWAS era. Existing databases like Pancan-meQTL and ASMdb have contributed to uncovering the role of genetic variants in cancer by linking DNA variations to methylation. However, they lack comprehensive regulatory annotations for such sites and rely solely on genomic proximity for target gene prediction, overlooking distal interactions. Our analysis using 3D interaction data reveals that over 94.8% of ASMs exhibit distal regulatory relationships, emphasizing their critical role in cancer epigenetics. Furthermore, these databases cover a limited number of cancer types (Pancan-meQTL: 23, ASMdb: 8). To address these limitations, we developed CanASM, a database that provides ASMs identified from BS-seq data for 31 human cancers, along with multidimensional regulatory annotations for ASM sites. This includes target genes, cis-regulatory elements (enhancers, ATAC-seq regions, insulators, 3D interactions, CpG islands), TFBSs, and GWAS cancer trait associations. Additionally, CanASM offers three interactive tools for analyzing TF binding affinity changes, predicting ASM target genes through 3D interactions, and identifying cell marker variations. As a comprehensive resource for SNV-driven epigenetic regulation in cancer, CanASM is helpful for in-depth research in cancer genetics and epigenetics.

## Introduction

Substantial evidence has confirmed that single-nucleotide variations (SNVs), particularly those within gene promoters, extensively influence DNA methylation levels at specific CpG sites. This knowledge contributes to our understanding of the complex regulatory mechanisms underlying the expression of specific genes and their roles in cancer. For example, the T allele at rs1001179 in the CAT promoter increases the levels of ETS-1 and GR-β transcription factors (TFs) by altering promoter methylation at specific CpGs and leading to increased CAT expression in chronic lymphocytic leukemia (CLL), which is associated with a more aggressive clinical course (1). Similarly, the core promoter of TERT, one of the most frequently mutated noncoding cancer drivers, exhibits high DNA methylation in Hep-G2 cells, effectively silencing its expression in a mutation-dependent manner (2). Another study demonstrated that rs7247241, located within the PPP1R14A promoter, exhibits allele-specific methylation, with the T allele showing increased CpG methylation and the C allele exhibiting stronger binding to CTCF and PLAGL2, thereby regulating PPP1R14A transcription (3). Despite these insights, many SNVs have not been linked to DNA methylation changes in different cancer types, and the effects of SNVs outside promoter regions on DNA methylation require further investigation.

Recent studies have shown that the SNVs that significantly affect CpG methylation are highly enriched within enhancer regions (4-8). In addition to enhancers and promoters, functional SNVs or CpG sites in multiple types of cis-regulatory elements have been identified. For example, rs6854845, located within a super enhancer (SE) region, disrupts long-range chromatin interactions with downstream target genes, potentially contributing to colon cancer progression, as demonstrated by chromatin conformation capture (3C) and real-time PCR analysis (9). Similarly, an assay for transposase-accessible chromatin sequencing (ATAC-seq) analysis during human preimplantation development revealed that open chromatin regions overlap extensively with hypomethylated distal elements, indicating their regulatory potential beyond CpG-rich promoters (10). 3D genome studies further suggest that CpG methylation prevents CTCF binding, disrupting chromatin loops that facilitate SNP−target gene cross-talk, whereas low CpG methylation increases loop formation, acting as an insulator to block long-range regulation in prostate cancer (4, 11). These findings highlight the need for systematic identification of SNV−CpG interactions across multiple regulatory elements of genes, especially those in distal regions, to gain deeper insights into transcriptional regulation in cancer.

Several high-throughput methods, including BeadChip-based methylation quantitative trait locus (meQTL) analysis (12-14) and the read count-based ASM approach (7, 15-17), have been developed to explore SNV−DNA methylation associations. The ASM approach uses bisulfite sequencing (BS-Seq) data to identify differences in CpG methylation between alleles of adjacent heterozygous SNV sites in homologous genomes, providing an effective strategy for linking genetic and epigenetic changes. Currently, two major databases have been established to catalog and provide insights into SNV−CpG relationships: Pancan-meQTL, which provides meQTLs for 23 cancer types in The Cancer Genome Atlas (TCGA) by integrating genotype and DNA methylation data (18), and ASMdb, which compiles ASM results across multiple species, including human cancer samples from eight tissue types (15). While these resources provide valuable insights into genetic and epigenetic interactions in cancer, no existing database currently provides ASM identification alongside cancer-specific regulatory annotations. From another perspective, the growing number of studies exploring the relationships between SNVs and DNA methylation, along with the rapid expansion of distinct cis-regulatory elements, TF ChIP-seq binding site (TFBS) and BS-seq datasets for human cancers, has prompted an urgent need to systematically identify ASM events and compile relevant cis-regulatory elements and TFs for a comprehensive exploration of ASM regulation across various cancer types.

To meet these needs, we developed CanASM, the first comprehensive database dedicated to genome-wide ASM identification and annotation across diverse cancers. CanASM provides a detailed map of ASM sites across 31 cancer types, along with annotations for the associated genes and regulatory elements, by integrating data for multiple cis-regulatory elements, TFBSs, TF binding affinity changes, etc. Here, cis-regulatory elements include typical enhancers, SEs, ATAC regions, 3D chromatin interaction anchors and CpG islands (CGIs). Users can search for ASMs according to SNV, genomic region or gene name. Additionally, CanASM enables functional analyses, including TF binding affinity alteration, target gene prediction through 3D interactions and cell marker variation prediction. By integrating large-scale BS-seq data with advanced regulatory annotations, CanASM offers a valuable resource for investigating SNV-driven epigenetic regulation in cancer.

## Materials and methods

### BS-seq data

BS-Seq is a powerful technique for measuring cytosine methylation at cytosine−phosphate−guanine (CpG) sites on a genome-wide scale and can also be used to identify SNVs. To construct CanASM, we queried the NCBI GEO database for all available bisulfite sequencing datasets as of March 2024, excluding low-quality and noncancer datasets. Ultimately, 226 BS-Seq samples were incorporated into this study (Supplementary Table S1), covering 31 cancer types and their matched normal tissue samples across 30 distinct tissues. The primary sequencing strategy was whole-genome bisulfite sequencing (WGBS, 63%), with additional data from reduced representation bisulfite sequencing (RRBS) and Capture-BS, etc.

### Cis-regulatory elements

To annotate the identified ASMs, we compiled six key types of cis-regulatory elements.

- Super enhancer (SE): Retrieved from the SEanalysis 2.0 web tool, which identified SEs from 1,739 human H3K27ac ChIP-seq samples (19).
- Typical enhancers: Retrieved from EnhancerAtlas2.0 (containing 192,173 human enhancers from 277 tissues/cells) (31740966) and Fantoms5 (containing 63,286 human enhancers from 1,829 samples) (20).
- ATAC regions: Retrieved from ATACdb (1,400 ATAC-seq human samples) (21) and a TCGA-based study (410 tumor samples across 23 cancer types) (22).
- 3D chromatin interactions: Retrieved from ENCODE (284 human 3D chromatin structure samples) (23), HiChIPdb (200 high-throughput HiChIP samples across 108 cell types, 10 kb resolution) (24), 4D_Nucleome (interactions from compressed interaction mcool data for 377 samples, 10 kb resolution) (25), and Chromloops (107 unique samples after excluding duplicates from the other three sources) (26). Ultimately, approximately 121 million 3D chromatin interactions from a total of 966 samples were retained.
- Insulators: Defined as CCCTC-binding factor (CTCF) ChIP-seq peaks (27), retrieved from the ChIP-atlas 3.0 (28) and ENCODE (29) databases, encompassing a total of 2,636 samples.
- CGIs: Retrieved from the UCSC Genome Browser (30) and from previous studies (31).

The reference genome version for all elements was GRCh38; for source data with the hg19 version, liftOver software (32) downloaded from UCSC was used to convert the data to GRCh38.

### TF ChIP-seq peaks and recognition motifs

To annotate the identified ASMs to TFBSs and analyze changes in TF binding affinity caused by ASM, we collected TFBSs and TF motifs for various TFs. TFBSs (GRCh38) for 1,980 TFs across more than 30,000 samples were integrated from the ChIP-atlas 3.0 (28) and ENCODE (29) databases. The position frequency matrices for ∼8,000 motifs, corresponding to 1,171 TFs, were retrieved from the JASPAR (33) and HOCOMOCOv12 (34) databases.

### Cancer GWAS datasets

To investigate the associations between index SNVs and cancers from a different perspective, we incorporated high-confidence cancer genome-wide association study (GWAS) data. Over 200 cancer-related GWAS datasets linking SNVs to approximately 30 different cancer types (p value <=0.001) were obtained from the OpenGWAS database (https://gwas.mrcieu.ac.hk/) and the GWAS Catalog (35). OpenGWAS is a manually curated collection of GWAS resources, whereas the GWAS Catalog offers tens of thousands of variant–trait pairs derived from high-quality, curated publications.

### Cell marker genes

SNPs have been demonstrated to play crucial roles in discerning cellular heterogeneity in cancer (36). To investigate the relationships between index SNVs and distinct cell types, we mapped five categories of ASM-associated genes (adjacent genes, eQTL genes, CpG target genes, distal genes via 3D interactions and proximal genes via 3D interactions) to cell identity marker genes. These markers were obtained from the CellMarker 2.0 database, which includes 16,679 manually curated markers across 429 human tissues and 1,715 cell types (37).

### ASM identification

The analytical pipeline used to identify ASMs is diagrammed in Figure1. The BS-seq reads of a sample were trimmed and filtered to remove sequencing adapters (default parameters if not specified) in GEO, low-quality bases (Phred score < 30) and those shorter than 40 bp using (Version 0.6.10), https://www.bioinformatics.babraham.ac.uk/projects/trim_galore/). The trimmed reads were aligned to the human genome (GRCh38) using Bismark (version 0.23.1) (38). Sambamba was used to remove duplicates (markdup, -r) (39). SNVs and CpG methylation were called using BS-SNPer (40) and Bismark, respectively. BS-SNPer was developed to identify SNV sites from BS-Seq data with high sensitivity and specificity without relying on an existing SNP library. The minimum mutation read number was set to 5 to ensure accuracy when calling SNVs. Then, heterozygous SNVs (GT:0/1) with a BS-SNPer-reported quality value of ‘pass’ were selected for further analysis.

To explore the methylation imbalance between the reference and alternative alleles, reads were first assigned to the respective alleles. C/T substitutions caused by bisulfite conversion impacts read assignment at variant sites, except for A/T and T/A mutations. For example, C/T mutations on the positive strand cannot be distinguished from C/T substitutions caused by bisulfite conversion; therefore, only reads supporting C or T alleles on the negative strand were used for ASM identification. We developed a read assignment strategy for each of the 12 mutation types (Table 1), which is consistent with the SNV calling process. For an SNV−CpG pair, overlapping reads were used for ASM identification through Fisher’s exact test (Figure 1C).

**Table 1.**
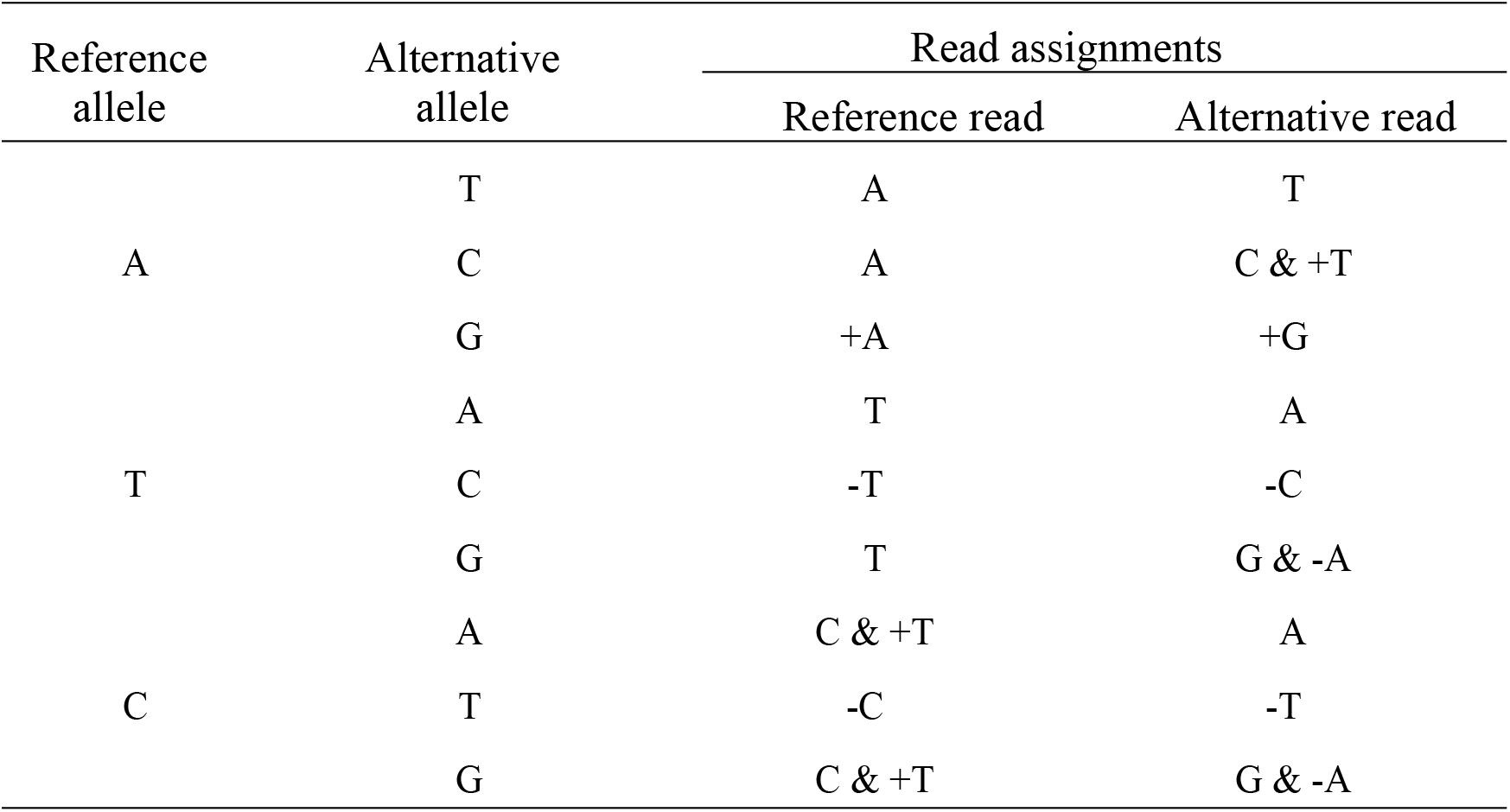
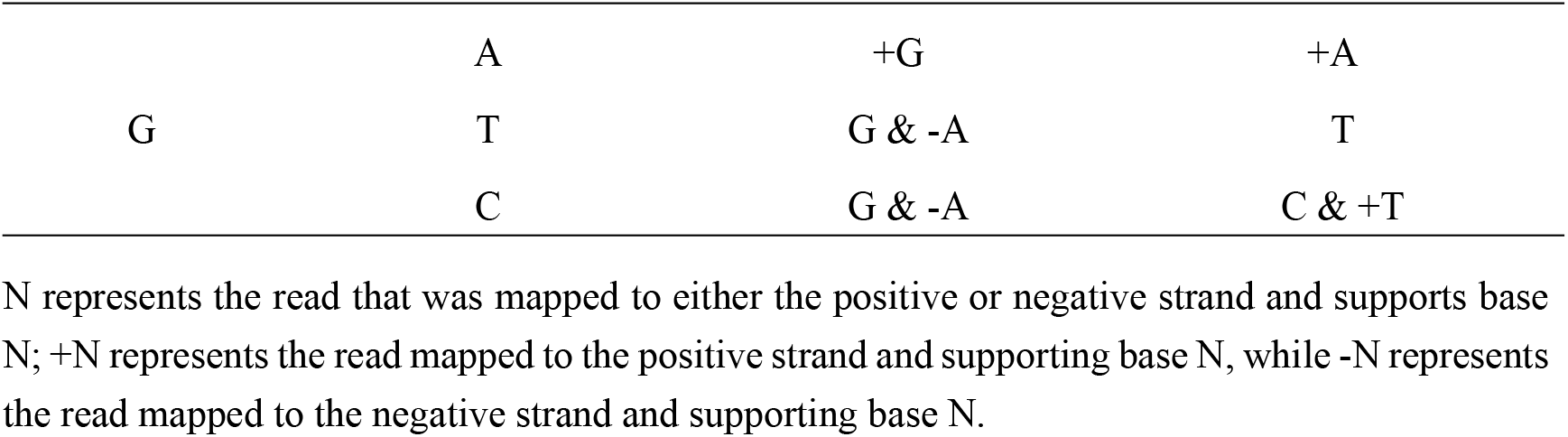
Allele assignments for each read covering SNV sites.

**Figure 1.**
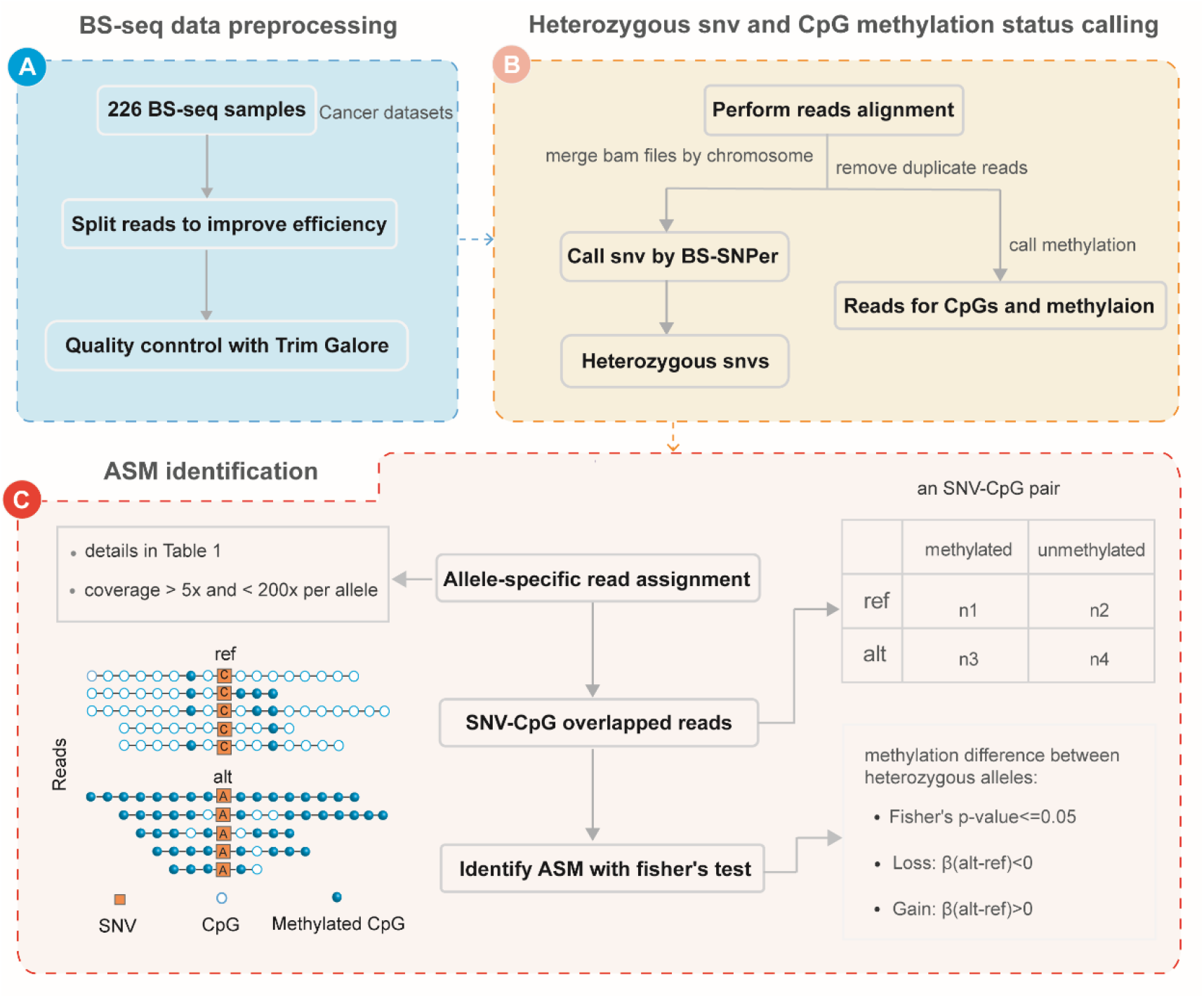
Analytical pipeline for ASM identification. A. The BS-seq data preprocessing process, including read splitting and quality control. B. Genotype of heterozygous SNV and CpG methylation status calling. C. Identification of ASM with Fisher’s exact test.

### SNV-to-SNP mapping

Since BS-SNPer identifies SNVs independently of known SNP datasets, the output includes only the genomic coordinates of SNVs. As a result, we initially assigned names to the SNVs on the basis of their position. If a SNP in the NCBI dbSNP database (v156; GRCh38 assembly) matched the position of an SNV, we renamed the SNV with the corresponding SNP name from the dbSNP database (41).

### Gene linkage to ASMs

To establish the associations between ASMs and their potential target genes, we implemented four distinct strategies as follows:

1. Genomic annotation. We utilized ANNOVAR (42) to annotate ASM index SNVs to the GRCh38 reference genes, with background genomic annotation files for reference genes downloaded from UCSC (http://hgdownload.cse.ucsc.edu/goldenPath/hg38/database/). This approach links ASMs to adjacent genes on the basis of their positions relative to the genes, including genes within 1 kb upstream/downstream regions, intergenic regions, exonic regions, intronic regions, and untranslated regions (UTRs).
2. Expression quantitative trait locus (eQTL) annotation. We obtained eQTLs of 49 tissues from the TargetGene database (43) to annotate ASM index SNVs.
3. CpG targeting. We retrieved the cancer-specific target genes for ASM-associated CpG sites from our previous study (44), which employs RNA-seq expression profiles and DNA methylation array data to computationally predict target genes of CpG sites across 33 different cancer types.
4. 3D chromatin interaction-based prediction. We leveraged chromatin interaction anchors to determine the spatial relationship between ASM index SNVs and gene promoters (defined as 2 kb upstream and downstream of transcription start sites). If an SNV overlapped with the same interaction anchor as a gene promoter, it was considered to have a proximal regulatory relationship with the gene. In contrast, when the SNV and gene promoter were located in separate interaction anchors within a 3D interaction pair, they were classified as having a distal regulatory relationship.

### Annotating ASMs to cis-elements and TFs

ASMs were annotated to the aforementioned distinct cis-regulatory elements and ChIP-seq peaks of TFs using BEDTools (45).

### Allele-specific binding analysis for TFs

Previous studies (6, 8) have supported the hypothesis that ASM correlates with allele-specific binding affinities of TF recognition motifs in regulatory elements. CanASM offers affinity alteration analysis at index SNVs for motifs using atSNP, which uses an importance sampling algorithm and a first-order Markov model to test the significance of SNP-driven changes in affinity scores (46).

## Results

### Content of CanASM

CanASM offers a comprehensive resource for ASM identification and the prediction of potential target genes, dissecting the associations between genetic variants and cancer at multiple molecular levels (Figure 2). The current version of CanASM catalogs 5,003,877 unique SNV–CpG pairs, including 3,056,776 distinct index SNVs and 4,157,508 CpGs, identified from 226 bisulfite BS-seq samples across 30 diverse human tissues. Of the total index SNVs, 2,634,406 (86%) were identified as SNPs. Importantly, CanASM provides detailed annotations of ASMs to a wealth of cis-regulatory elements and TF binding sites in detail. Furthermore, CanASM leverages 3D chromatin interaction data to predict proximal and distal target genes for ASMs, while also incorporating adjacent genes, CpG-associated target genes, and eQTLs as potential targets for ASMs (as detailed in the Methods), offering a multi-dimensional perspective on the prediction of ASM-regulated genes. In addition, CanASM also offer cancer GWAS annotation for index SNVs. Moreover, CanASM provides three online analysis tools, including allele-specific binding analysis for TFs, target genes prediction based on 3D interaction, and cell marker SNV prediction. Additionally, CanASM supports batch browsing and downloading of ASMs, enabling efficient access to large-scale data. Overall, CanASM mainly provides a user-friendly interface to query, analyze, browse and download of regulatory information and potential target genes for ASMs identified from cancer related samples.

**Figure 2.**
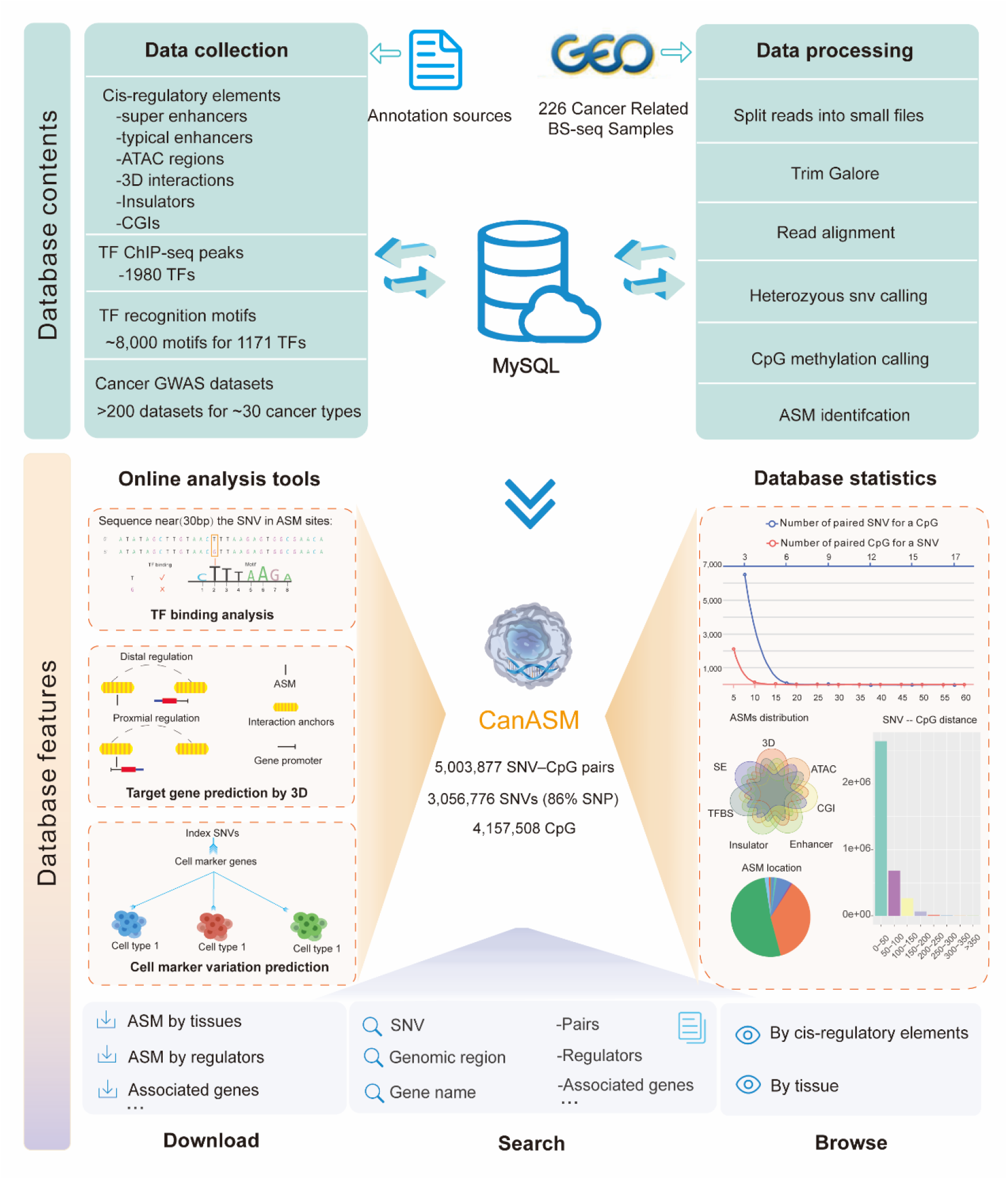
Schematic overview of CanASM. Overview of BS-seq data collection and processing, annotation data collection, and database features of CanASM.

### Feature and utility of CanASM

#### Database statistics

The results of mapping ASMs to regulatory elements indicate a widespread distribution of index SNVs across key regulatory elements, with 3,048,821 (99.7%) index SNVs located in 3D chromatin interaction regions, 2,721,528 (89.0%) in typical enhancers, 1,461,174 (47.8%) in super enhancers, 1,278,694 (41.8%) in ATAC-seq regions, and 1,707,588 (55.9%) in TFBSs. In addition, 314,096 (10.3%) index SNVs are found in insulators, whereas 214,630 (7.0%) target CpG sites within CpG islands. Interestingly, the target gene prediction results based on 3D interactions revealed that 2,898,618 index SNVs (94.8%) presented distal regulatory relationships with their target genes, whereas only 166,579 (5.2%) presented proximal regulatory relationships. This result highlights the crucial role of ASMs in mediating long-range chromatin interactions and gene regulation, offering novel insights into the spatial organization of the genome in transcriptional regulation.

Specifically, the statistical interface of CanASM includes statistics on the distribution of ASMs across gene contexts, distances between index SNVs and CpGs, methylation change patterns at ASM sites, ASMs across chromosomes, ASMs among regulatory elements and SNV−CpG pairs, offering an overview of ASM characterization (Figure 2).

#### Advanced search

CanASM has been designed with user-friendly web interfaces that make it easy to access detailed annotations for ASMs. Users can query ASMs by entering an SNV, genomic region or gene name on the home page (Figure 3A). When the “SNV” parameter is selected, users can input an SNV by specifying either the SNP name (e.g., rs5018177) if the SNV is included in the dbSNP database or the chromosomal position (e.g., chr1:10004334) if the SNV is not included in the dbSNP database. When the “genomic region” parameter is selected, all index SNVs within the specified region are returned. When the “associated gene” parameter is selected, all index SNVs linked to the gene according to the four SNV−gene association strategies (see Methods) are returned. Upon submission of the query, the results interface initially displays a list of ASM events, including SNV−CpG pairs, sample identifiers, and tissue types, associated with the queried SNVs across all available samples. The interface also enables users to dynamically filter ASMs according to the tissue of interest. Users can select a specific SNV of interest to navigate to a detailed view.

**Figure 3.**
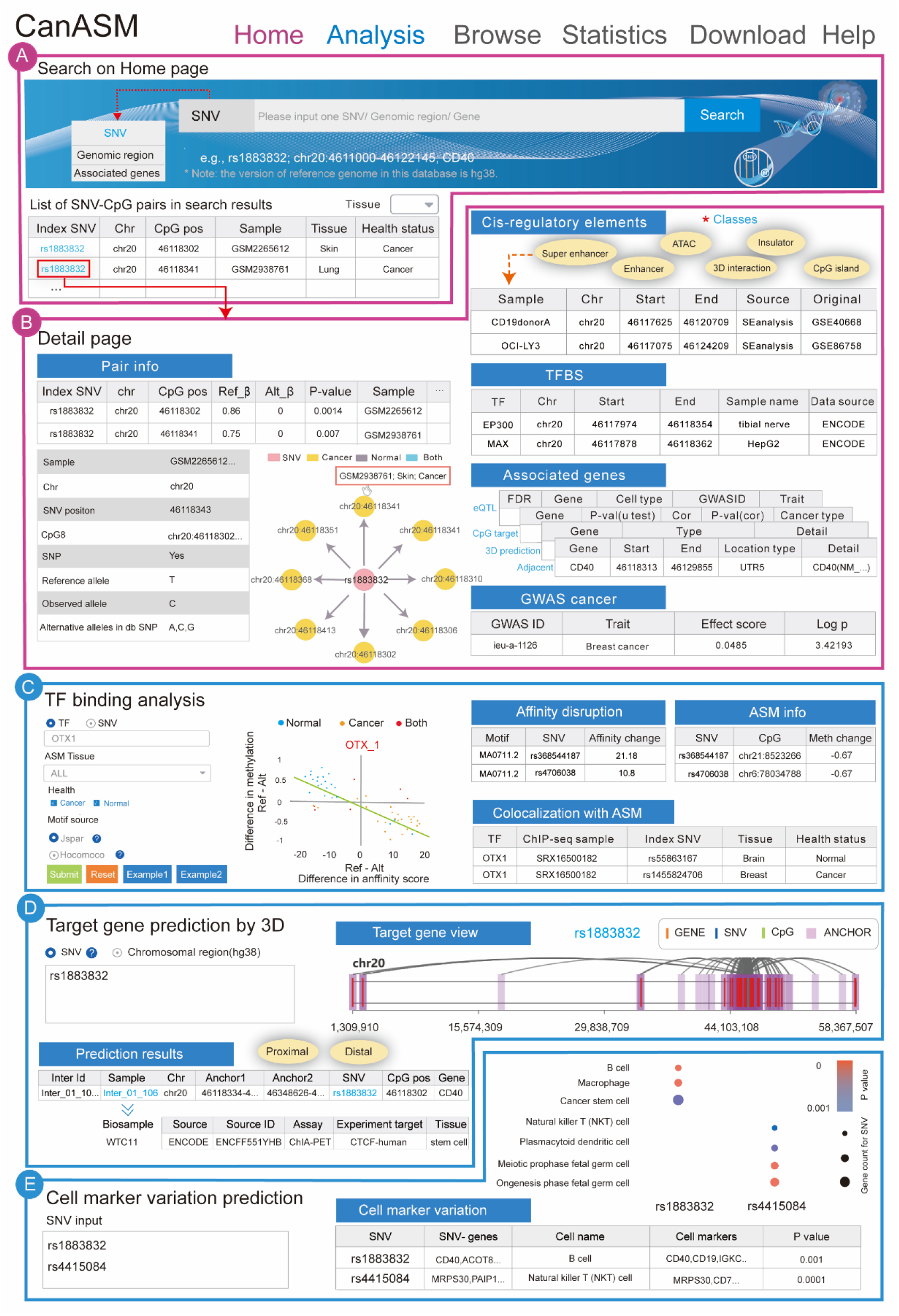
Database usage. A. The ‘Home’ page provides a global search function for for SNV–CpG pairs across all available samples though entering an SNV, gene or genomic regions. B. The detail page for an index SNV provides annotations of cis-regulatory elements, related genes, TFBSs and cancer GWAS traits. C. TF binding analysis provides TF binding affinity changes, the correlation between binding affinity alterations and methylation changes, and the TFBSs at ASM sites. D. The target gene prediction on the basis of 3D interactions provides relationships between the promoters of potential target genes and ASMs, including both distal and proximal regulation. E. Cell Marker SNVs analysis provides relationships between index SNVs and different cell identities through performing enrichment analysis using the hypergeometric test with the SNV-related gene sets and cell marker genes.

Upon selection of a specific SNV, users are directed to a detail interface that systematically organizes multidimensional annotations for the index SNV (Figure 3B). This interface includes details for CpG sites targeted by the SNV and associated cis-regulatory elements, including enhancers, super enhancers, ATAC-seq accessible regions, 3D chromatin interaction domains, insulators, and CpG islands (CGIs). Furthermore, related genes categorized as adjacent genes, 3D-predicted genes, eQTL genes, and CpG-targeted genes are also included. Additionally, the detailed interface displays TFBSs and cancer GWAS traits linked to the SNV.

#### Online analysis tools

CanASM provides three analysis tools, including tools for TF binding analysis, target gene prediction on the basis of 3D interactions and cell marker variation prediction, to further explore the underlying mechanisms of ASMs in an interactive manner.

TF binding analysis is designed to facilitate the investigation of regulatory effects induced by ASMs through the analysis of TF binding affinity changes, the correlation between binding affinity alterations and methylation changes, and the identification of ASMs located within TFBSs (Figure 3C). Users can input a specific TF name to evaluate changes in its binding affinity caused by index SNVs using PFMs of motifs or input a specific SNV to identify all the affected TFs. Notably, if a TF is input, the tool also provides an analysis of the correlation between changes in binding affinity and methylation levels of paired CpGs influenced by ASMs across all samples. To increase flexibility, users can select background ASMs according to sample types and tissue origins of ASMs and select different motif data sources to optimize the prediction TF binding affinity changes and colocalization with TFBSs. This tool offers computational support for exploring the relationships between TF binding dynamics at ASM sites.

The tool for target gene prediction on the basis of 3D interactions is designed to predict the target genes for ASMs by analyzing the positional relationships among ASM sites, gene promoters, and the two anchor points of 3D chromatin interaction pairs (Figure 3D). In the case of distal regulation, the index SNV is located at one anchor point of a chromatin interaction pair, and the promoter of the target gene resides at the other anchor point, indicating that the regulatory influence of the ASM occurs through long-range chromatin looping. Conversely, proximal regulation is characterized by the ASM site situated within the same anchor point as the gene promoter. To increase flexibility, users can filter background ASMs on the basis of sample tissues. This tool enables researchers to utilize large-scale 3D chromatin interaction data to predict target genes for genomic regions or SNVs of interest. If users input genomic regions, all index SNVs located within those regions are analyzed.

Cell marker variation prediction is designed to predict cell types marked by genes associated with ASMs from user-specified SNVs (Figure 3E). Users can input SNVs of interest in one of two standardized formats: a reference SNP ID from dbSNP (e.g., rs1883832) or a chromosomal position (e.g., chr1:1548485) if the SNV is not recorded in dbSNP. We performed enrichment analysis using the hypergeometric test to assess the association between SNV-related gene sets and cell marker genes, aiming to identify Cell Marker SNVs. To increase flexibility, users can filter background ASMs on the basis of the sample tissue. This tool facilitates the identification of cell identities mediated by ASM-associated genetic variations.

#### Data download

All ASM (i.e., SNV−CpG pairs) information is categorized by tissue and available for download. Additionally, the maps of index SNVs and various cis-regulatory elements (including super enhancers, typical enhancers, ATAC regions, 3D interaction regions, and CpG islands), the maps of index SNVs and TFBSs, the relationships between index SNVs and motif disruptions, and the links between index SNVs and GWAS cancer associations are all provided for download.

#### Data browse

All ASM data, categorized by tissue and various cis-elements, can be interactively browsed in batches.

### Case study

CanASM has a wide range of application scenarios. One of the challenges in the post-GWAS era is identifying target genes and elucidating the regulatory mechanisms of cancer-associated variants. Here, we demonstrated how to achieve this goal with CanASM using rs1883832, a recently identified lung cancer risk SNP, as an example. A prior study used MassARRAY to genotype six candidate SNPs in a cohort of 400 lung cancer patients and 400 healthy controls, revealing a significant association between rs1883832 and increased lung cancer risk (P < 0.001) (47). However, further research is needed to elucidate the molecular function of rs1883832 in lung cancer. To this end, we used CanASM to systematically examine the regulatory effects of rs1883832.

We searched rs1883832 in CanASM and identified six CpG sites in skin and lung cancer tissues that display allele-specific methylation at rs1883832. These six CpG sites are located within 40 bp of rs1883832, and two of these sites are found in lung cancer, exhibiting hypermethylation near the mutant allele (C). The detailed page of the cis-regulatory element module shows that rs1883832 overlaps with cis-regulatory elements, including super enhancers, ATAC regions, insulators, and 3D chromatin interaction regions. Additionally, its targeted CpG sites are situated within CpG islands. These findings emphasize the potential regulatory significance of the target genes. Furthermore, the TF binding module revealed that rs1883832 is located within the ChIP-seq binding sites of more than 70 transcription factors, including MYC and TP53BP1. Notably, MYC is a well-established oncogene that plays pivotal roles in cell proliferation, apoptosis, and cancer progression (48-51), and TP53BP1 is a critical tumor suppressor involved in DNA damage repair and maintenance of genomic stability (52-55) . Moreover, the associated gene module shows that rs1883832 is in the UTR5 region of the tumor necrosis factor receptor CD40, a novel immunomodulatory cancer therapy target (56).

With the online target gene prediction tool, we identified extensive 3D chromatin interaction data suggesting a potential proximal regulatory relationship between rs1883832 and CD40 (where the promoter and rs1883832 were located within the same interaction anchor). Additionally, rs1883832 exhibited potential distal regulatory relationships (where the promoter and rs1883832 are located at opposite ends of interaction anchors) with more than sixty genes, including MMP9 and UBE2C. Notably, MMP9 is a key enzyme involved in cancer progression, invasion, and metastasis due to its role in degrading basement membranes (57, 58), whereas UBE2C, a ubiquitin-conjugating enzyme, promotes cancer progression primarily through its role in the ubiquitin−proteasome system, facilitating the degradation of key cell cycle regulators in multiple cancers (59-61). Furthermore, the cell marker variation online tool revealed that rs1883832 is a marker variation for many cancer cells, such as “cancer stem cells” and “M2 macrophages”.

These findings highlight the potential role of ASM in epigenetic regulation and tumor progression and demonstrate that CanASM is a valuable resource for researchers investigating cancer-associated genetic variation and epigenetic regulation.

## Discussion

The identification of ASM is crucial for understanding the interplay between genetic variations and epigenetic regulation in cancer. While GWASs have successfully identified numerous cancer-associated SNVs, the precise mechanisms by which these variants influence gene regulation remain largely unresolved. ASM has emerged as a key regulatory mechanism, particularly when it occurs within enhancers and other cis-regulatory elements, where genetic variations can lead to differential methylation patterns that alter gene expression. Existing databases, such as Pancan-meQTL and ASMdb, provide significant insights into the associations between DNA variations and methylation status. However, these resources lack comprehensive regulatory annotations, such as information on cis-regulatory elements and TFBSs. Furthermore, their methodologies for target gene prediction rely solely on genomic proximity, failing to account for the complexities of distal regulatory interactions that may play a critical role in gene regulation. Our results indicate that more than 94.8% of the index SNVs for ASMs exhibit potential distal regulatory relationships with predicted target genes, emphasizing the important role of ASM in spatial genome organization in cancer epigenetics. Additionally, these databases cover a limited number of cancer types, with Pancan-meQTL encompassing 23 types and ASMdb including 8 types. Given the rapid accumulation of BS-seq datasets and the substantial growth of diverse regulators, a systematic database dedicated to ASM identification and annotation for cancer is needed. CanASM was developed to fill this gap, offering a comprehensive resource for genome-wide ASM identification and functional annotation across diverse cancer types.

We developed a customized computational pipeline (detailed in the Methods section) to detect allele-specific methylation using BS-seq data, constructing a comprehensive ASM landscape for human cancer. Notably, beyond ASM detection, CanASM offers multidimensional regulatory annotations for ASMs through systematic integration of putative target genes; cis-regulatory elements, including typical enhancers, super enhancers, ATAC-seq regions, insulators, 3D interaction regions and CGIs; TFBSs; and GWAS cancer trait associations. To further increase its utility, CanASM incorporates three interactive online analysis tools, enabling users to find TF binding affinity changes at ASM sites, predict ASM target genes through 3D chromatin interactions, and investigate cell marker variations associated with ASMs. In addition, CanASM supports statistical visualization, interactive browsing, and bulk data retrieval. Despite its strengths, CanASM has certain limitations that require further optimization. While CanASM leverages extensive publicly available bisulfite sequencing datasets, the database is constrained by the sample diversity of existing studies. Future updates incorporating single-cell BS-seq data and long-read sequencing technologies could increase the resolution of ASM mapping, enabling more precise identification of cell type-specific regulatory mechanisms and improving our understanding of genetic−epigenetic interactions in cancer.

In summary, CanASM provides the first comprehensive, cancer-specific database for ASM identification and functional annotation, integrating diverse regulatory datasets to facilitate the study of SNV-driven epigenetic regulation. By offering a systematic framework for ASM analysis, CanASM has the potential to accelerate the discovery of novel cancer biomarkers, regulatory mechanisms, and therapeutic targets.

## Funding information and Acknowledgments

This work was supported by the National Natural Science Foundation of China (Grant Nos. 62225109, 62402345, and 62302342). We would like to express our sincere thanks to the funding agencies for their financial support, which made this research possible.

## References

1. Galasso M, Dalla Pozza E, Chignola R, Gambino S, Cavallini C, Quaglia FM, et al. The rs1001179 SNP and CpG methylation regulate catalase expression in chronic lymphocytic leukemia. Cell Mol Life Sci. 2022;79(10):521.

2. Kouroukli AG, Rajaram N, Bashtrykov P, Kretzmer H, Siebert R, Jeltsch A, et al. Targeting oncogenic TERT promoter variants by allele-specific epigenome editing. Clin Epigenetics. 2023;15(1):183.

3. Tian Y, Soupir A, Liu Q, Wu L, Huang CC, Park JY, et al. Novel role of prostate cancer risk variant rs7247241 on PPP1R14A isoform transition through allelic TF binding and CpG methylation. Hum Mol Genet. 2022;31(10):1610–21.

4. Zeng Y, Jain R, Lam M, Ahmed M, Guo H, Xu W, et al. DNA methylation modulated genetic variant effect on gene transcriptional regulation. Genome Biol. 2023;24(1):285.

5. Villicana S, Castillo-Fernandez J, Hannon E, Christiansen C, Tsai PC, Maddock J, et al. Genetic impacts on DNA methylation help elucidate regulatory genomic processes. Genome Biol. 2023;24(1):176.

6. Do C, Dumont ELP, Salas M, Castano A, Mujahed H, Maldonado L, et al. Allele-specific DNA methylation is increased in cancers and its dense mapping in normal plus neoplastic cells increases the yield of disease-associated regulatory SNPs. Genome Biol. 2020;21(1):153.

7. Onuchic V, Lurie E, Carrero I, Pawliczek P, Patel RY, Rozowsky J, et al. Allele-specific epigenome maps reveal sequence-dependent stochastic switching at regulatory loci. Science. 2018;361(6409).

8. Stefansson OA, Sigurpalsdottir BD, Rognvaldsson S, Halldorsson GH, Juliusson K, Sveinbjornsson G, et al. The correlation between CpG methylation and gene expression is driven by sequence variants. Nat Genet. 2024;56(8):1624–31.

9. Cong Z, Li Q, Yang Y, Guo X, Cui L, You T. The SNP of rs6854845 suppresses transcription via the DNA looping structure alteration of super-enhancer in colon cells. Biochem Biophys Res Commun. 2019;514(3):734–41.

10. Wu J, Xu J, Liu B, Yao G, Wang P, Lin Z, et al. Chromatin analysis in human early development reveals epigenetic transition during ZGA. Nature. 2018;557(7704):256–60.

11. Ahmed M, Soares F, Xia JH, Yang Y, Li J, Guo H, et al. CRISPRi screens reveal a DNA methylation-mediated 3D genome dependent causal mechanism in prostate cancer. Nat Commun. 2021;12(1):1781.

12. Hawe JS, Wilson R, Schmid KT, Zhou L, Lakshmanan LN, Lehne BC, et al. Genetic variation influencing DNA methylation provides insights into molecular mechanisms regulating genomic function. Nat Genet. 2022;54(1):18–29.

13. Meeks GL, Henn BM, Gopalan S. Genetic differentiation at probe SNPs leads to spurious results in meQTL discovery. Commun Biol. 2023;6(1):1295.

14. Cheng Y, Li B, Zhang X, Aouizerat BE, Zhao H, Xu K. Reply to: Genetic differentiation at probe SNPs leads to spurious results in meQTL discovery. Commun Biol. 2023;6(1):1296.

15. Zhou Q, Guan P, Zhu Z, Cheng S, Zhou C, Wang H, et al. ASMdb: a comprehensive database for allele-specific DNA methylation in diverse organisms. Nucleic Acids Res. 2022;50(D1):D60–D71.

16. Dumont ELP, Tycko B, Do C. CloudASM: an ultra-efficient cloud-based pipeline for mapping allele-specific DNA methylation. Bioinformatics. 2020;36(11):3558–60.

17. Rosenski J, Peretz A, Magenheim J, Loyfer N, Shemer R, Glaser B, et al. Atlas of imprinted and allele-specific DNA methylation in the human body. Nat Commun. 2025;16(1):2141.

18. Gong J, Wan H, Mei S, Ruan H, Zhang Z, Liu C, et al. Pancan-meQTL: a database to systematically evaluate the effects of genetic variants on methylation in human cancer. Nucleic Acids Res. 2019;47(D1):D1066–D72.

19. Qian FC, Zhou LW, Li YY, Yu ZM, Li LD, Wang YZ, et al. SEanalysis 2.0: a comprehensive super-enhancer regulatory network analysis tool for human and mouse. Nucleic Acids Res. 2023;51(W1):W520-W7.

20. Noguchi S, Arakawa T, Fukuda S, Furuno M, Hasegawa A, Hori F, et al. FANTOM5 CAGE profiles of human and mouse samples. Sci Data. 2017;4:170112.

21. Wang F, Bai X, Wang Y, Jiang Y, Ai B, Zhang Y, et al. ATACdb: a comprehensive human chromatin accessibility database. Nucleic Acids Res. 2021;49(D1):D55–D64.

22. Dahlin AM, Wibom C, Ghasimi S, Brannstrom T, Andersson U, Melin B. Relation between Established Glioma Risk Variants and DNA Methylation in the Tumor. PLoS One. 2016;11(10):e0163067.

23. Consortium EP. An integrated encyclopedia of DNA elements in the human genome. Nature. 2012;489(7414):57–74.

24. Zeng W, Liu Q, Yin Q, Jiang R, Wong WH. HiChIPdb: a comprehensive database of HiChIP regulatory interactions. Nucleic Acids Res. 2023;51(D1):D159–D66.

25. Dekker J, Belmont AS, Guttman M, Leshyk VO, Lis JT, Lomvardas S, et al. The 4D nucleome project. Nature. 2017;549(7671):219–26.

26. Zhou Q, Cheng S, Zheng S, Wang Z, Guan P, Zhu Z, et al. ChromLoops: a comprehensive database for specific protein-mediated chromatin loops in diverse organisms. Nucleic Acids Res. 2023;51(D1):D57–D69.

27. Chen W, Zeng YC, Achinger-Kawecka J, Campbell E, Jones AK, Stewart AG, et al. Machine learning enables pan-cancer identification of mutational hotspots at persistent CTCF binding sites. Nucleic Acids Res. 2024;52(14):8086–99.

28. Zou Z, Ohta T, Oki S. ChIP-Atlas 3.0: a data-mining suite to explore chromosome architecture together with large-scale regulome data. Nucleic Acids Res. 2024;52(W1):W45-W53.

29. Davis CA, Hitz BC, Sloan CA, Chan ET, Davidson JM, Gabdank I, et al. The Encyclopedia of DNA elements (ENCODE): data portal update. Nucleic Acids Res. 2018;46(D1):D794–D801.

30. Raney BJ, Barber GP, Benet-Pages A, Casper J, Clawson H, Cline MS, et al. The UCSC Genome Browser database: 2024 update. Nucleic Acids Res. 2024;52(D1):D1082–D8.

31. Su J, Zhang Y, Lv J, Liu H, Tang X, Wang F, et al. CpG_MI: a novel approach for identifying functional CpG islands in mammalian genomes. Nucleic Acids Res. 2010;38(1):e6.

32. Zhou T, Pu SY, Zhang SJ, Zhou QJ, Zeng M, Lu JS, et al. Dog10K: an integrated Dog10K database summarizing canine multi-omics. Nucleic Acids Res. 2025;53(D1):D939–D47.

33. Rauluseviciute I, Riudavets-Puig R, Blanc-Mathieu R, Castro-Mondragon JA, Ferenc K, Kumar V, et al. JASPAR 2024: 20th anniversary of the open-access database of transcription factor binding profiles. Nucleic Acids Res. 2024;52(D1):D174–D82.

34. Vorontsov IE, Eliseeva IA, Zinkevich A, Nikonov M, Abramov S, Boytsov A, et al. HOCOMOCO in 2024: a rebuild of the curated collection of binding models for human and mouse transcription factors. Nucleic Acids Res. 2024;52(D1):D154–D63.

35. Sollis E, Mosaku A, Abid A, Buniello A, Cerezo M, Gil L, et al. The NHGRI-EBI GWAS Catalog: knowledgebase and deposition resource. Nucleic Acids Res. 2023;51(D1):D977–D85.

36. Zhou Y, Zhao L, Cai M, Luo D, Pang Y, Chen J, et al. Utilizing sc-linker to integrate single-cell RNA sequencing and human genetics to identify cell types and driver genes associated with non-small cell lung cancer. BMC Cancer. 2025;25(1):130.

37. Hu C, Li T, Xu Y, Zhang X, Li F, Bai J, et al. CellMarker 2.0: an updated database of manually curated cell markers in human/mouse and web tools based on scRNA-seq data. Nucleic Acids Res. 2023;51(D1):D870–D6.

38. Krueger F, Andrews SR. Bismark: a flexible aligner and methylation caller for Bisulfite-Seq applications. Bioinformatics. 2011;27(11):1571–2.

39. Tarasov A, Vilella AJ, Cuppen E, Nijman IJ, Prins P. Sambamba: fast processing of NGS alignment formats. Bioinformatics. 2015;31(12):2032–4.

40. Gao S, Zou D, Mao L, Liu H, Song P, Chen Y, et al. BS-SNPer: SNP calling in bisulfite-seq data. Bioinformatics. 2015;31(24):4006–8.

41. Phan L, Zhang H, Wang Q, Villamarin R, Hefferon T, Ramanathan A, et al. The evolution of dbSNP: 25 years of impact in genomic research. Nucleic Acids Res. 2025;53(D1):D925–D31.

42. Wang K, Li M, Hakonarson H. ANNOVAR: functional annotation of genetic variants from high-throughput sequencing data. Nucleic Acids Res. 2010;38(16):e164.

43. Correction to ‘TargetGene: a comprehensive database of cell-type-specific target genes for genetic variants’. Nucleic Acids Res. 2024;52(D1):D1700.

44. Zhao J, Qian F, Li X, Yu Z, Zhu J, Yu R, et al. CanMethdb: a database for genome-wide DNA methylation annotation in cancers. Bioinformatics. 2023;39(1).

45. Quinlan AR, Hall IM. BEDTools: a flexible suite of utilities for comparing genomic features. Bioinformatics. 2010;26(6):841–2.

46. Zuo C, Shin S, Keles S. atSNP: transcription factor binding affinity testing for regulatory SNP detection. Bioinformatics. 2015;31(20):3353–5.

47. He J, Yu L, Qiao Z, Yu B, Liu Y, Ren H. Genetic polymorphisms of FCGR2A, ORAI1 and CD40 are associated with risk of lung cancer. Eur J Cancer Prev. 2022;31(1):7–13.

48. Chen J, Zhang S, Huang X, Wang Q, Xu W, Huang J, et al. Sialylated IgG-activated integrin beta4-Src-Erk axis stabilizes c-Myc in a p300 lysine acetyltransferase-dependent manner to promote colorectal cancer liver metastasis. Neoplasia. 2025;61:101140.

49. Zheng B, Wang YX, Wu ZY, Li XW, Qin LQ, Chen NY, et al. Design, Synthesis and Bioactive Evaluation of Topo I/c-MYC Dual Inhibitors to Inhibit Oral Cancer via Regulating the PI3K/AKT/NF-kappaB Signaling Pathway. Molecules. 2025;30(4).

50. Matos A, Ovens AJ, Jakobsen E, Iglesias-Gato D, Bech JM, Friis S, et al. Salicylate-Elicited Activation of AMP-Activated Protein Kinase Directly Triggers Degradation of C-Myc in Colorectal Cancer Cells. Cells. 2025;14(4).

51. Garcia Garcia A, Ferrer Aporta M, Vallejo Palma G, Giraldez Trujillo A, Castillo-Gonzalez R, Calzon Lozano D, et al. Targeting ELOVL6 to disrupt c-MYC driven lipid metabolism in pancreatic cancer enhances chemosensitivity. Nat Commun. 2025;16(1):1694.

52. Harvey-Jones E, Raghunandan M, Robbez-Masson L, Magraner-Pardo L, Alaguthurai T, Yablonovitch A, et al. Longitudinal profiling identifies co-occurring BRCA1/2 reversions, TP53BP1, RIF1 and PAXIP1 mutations in PARP inhibitor-resistant advanced breast cancer. Ann Oncol. 2024;35(4):364–80.

53. Georgieva D, Wang N, Taglialatela A, Jerabek S, Reczek CR, Lim PX, et al. BRCA1 and 53BP1 regulate reprogramming efficiency by mediating DNA repair pathway choice at replication-associated double-strand breaks. Cell Rep. 2024;43(4):114006.

54. Dibitetto D, Liptay M, Vivalda F, Dogan H, Gogola E, Gonzalez Fernandez M, et al. H2AX promotes replication fork degradation and chemosensitivity in BRCA-deficient tumours. Nat Commun. 2024;15(1):4430.

55. Kilgas S, Syed A, Toolan-Kerr P, Swift ML, Roychoudhury S, Sarkar A, et al. NEAT1 modulates the TIRR/53BP1 complex to maintain genome integrity. Nat Commun. 2024;15(1):8438.

56. Jian CZ, Lin L, Hsu CL, Chen YH, Hsu C, Tan CT, et al. A potential novel cancer immunotherapy: Agonistic anti-CD40 antibodies. Drug Discov Today. 2024;29(3):103893.

57. Yang Y, Song S, Li S, Kang J, Li Y, Zhao N, et al. GATA4 regulates the transcription of MMP9 to suppress the invasion and migration of breast cancer cells via HDAC1-mediated p65 deacetylation. Cell Death Dis. 2024;15(4):289.

58. Rademaekers M, Johansson EO, Johansson E, Roberg K, Wiechec E. Tumor-matched and unmatched cancer associated fibroblasts exhibit differential effect on proliferation and FMOD and MMP9 gene expression in head and neck squamous cell carcinoma cells when cocultured in spheroids. Cancer Cell Int. 2024;24(1):190.

59. Zhao M, Li J, Wang R, Mi L, Gu Y, Chen R, et al. Ubiquitination-Binding Enzyme 2C is Associated with Cancer Development and Prognosis and is a Potential Therapeutic Target. Onco Targets Ther. 2024;17:1159–71.

60. Guo Y, Chen X, Zhang X, Hu X. UBE2S and UBE2C confer a poor prognosis to breast cancer via downregulation of Numb. Front Oncol. 2023;13:992233.

61. Chen Q, Mo S, Zhu L, Tang M, Cheng J, Ye P, et al. Prognostic implication of UBE2C + CD8 + T cell in neoadjuvant immune checkpoint blockade plus chemotherapy for locally advanced esophageal cancer. Int Immunopharmacol. 2024;130:111696.

